# Neurotensin and Dynorphin Bi-Directionally Modulates CeA Inhibition of oval BNST Neurons in Male Mice

**DOI:** 10.1101/367649

**Authors:** CP Normandeau, ML Torruella Suárez, P Sarret, ZA McElligott, EC Dumont

## Abstract

Neuropeptides are often co-expressed in neurons but their neurophysiological effects are commonly studied individually. Multiple neuropeptides may therefore be simultaneously released to coordinate proper neural circuit function. Here, we triggered the release of endogenous neuropeptides in brain slices from male mice to better understand the modulation of central amygdala (CeA) inhibitory inputs onto oval (ov) BNST neurons. We found that locally-released neurotensin (NT) and dynorphin (Dyn) antagonistically regulated CeA inhibitory inputs onto ovBNST neurons. NT and Dyn respectively increased and decreased CeA-to-ovBNST inhibitory inputs through NT receptor 1 (NTR1) and kappa opioid receptor (KOR). Additionally, NT and Dyn mRNAs were highly co-localized in ovBNST neurons suggesting that they may be released from the same cells. Together, we showed that NT and Dyn are key modulators of CeA inputs to ovBNST, paving the way to determine whether different conditions or states can alter the neuropeptidergic regulation of this particular brain circuit.

## Introduction

Neuropeptides are frequently co-expressed in individual neurons within the nervous system; their cooperative role may enable flexible neuromodulation and proper function of neural circuits (Griebel and Holsboer, 2012; Kormos and Gaszner, 2013; Valentino and Aston-Jones, 2010). However, neuropeptides are often studied individually, neglecting the functional outcomes resulting from their coordinated actions (Ptak *et al*, 2009; Sun *et al*, 2003). In brain slices prepared from male rats, post-synaptic activation of oval bed nucleus of the stria terminalis (ovBNST) neurons triggers the release of various neuropeptides, such as neurotensin (NT), corticotrophin releasing factor (CRF) and dynorphin (Dyn) which in turn robustly modulate excitatory and inhibitory synaptic transmission onto the same neurons (Normandeau *et al*, 2018). Notably, endogenously-released NT produces an increase in inhibitory transmission in the rat ovBNST, that is further enhanced by chronic stress (Normandeau *et al*, 2018). What remains unknown is whether this coordinated modulation of synaptic transmission is circuit-specific.

Although still under debate, the ovBNST may be devoted to energy homeostasis, promoting foraging behaviours and food intake (Jennings *et al*, 2013; Li and Kirouac, 2008; Moga *et al*, 1995). The ovBNST is robustly and reciprocally connected with the central nucleus of the amygdala (CeA) and because these bi-directional connections are exclusively GABAergic, the ovBNST and the CeA most likely inhibit each other (Petrovich and Swanson, 1997). This connection may be integral in the balance between aversive or appetitive behaviours (Davis *et al*, 2010; Dong *et al*, 2001; Jennings *et al*, 2013). Whether neuropeptides are important modulators of this particular brain circuit is largely undetermined. Nonetheless, it is known that the opiate dynorphin (Dyn) released from GABA neurons specifically decreases CeA-to-dorsal BNST inhibitory inputs in male mice (Crowley *et al*, 2016; Li *et al*, 2012). Whether Dyn cooperates with other neuropeptides in the ovBNST to regulate CeA inhibition is currently unknown.

Using brain slice electrophysiology in male mice, we triggered endogenous release of neuropeptides and discovered that release of NT and Dyn opposingly modulated CeA-to-ovBNST synaptic inhibition through neurotensin receptors 1 (NTR1) and kappa opioid receptors (KOR), respectively. NT-mediated enhancement of inhibitory transmission in the ovBNST overshadowed the effect of Dyn, paving the way to determine whether changes in condition/state may affect neuropeptidergic regulation of synaptic transmission in this particular neural circuit.

## Materials and Methods

### Mice

46 mice (>8 weeks old) were group housed on a reverse 12-hour light/dark cycle (lights OFF at 8:00 A.M) with *ad libitum* access to chow and water. C57BL/6J adult male mice (n=29) used for electrophysiology were obtained from Charles River Laboratories (St-Constant, QC, Canada) and V*gat-Cre* adult male mice (n=17) used for electrophysiology and *in situ* hybridization were bred in house in the McElligott lab (University of North Carolina, Chapel Hill, NC, USA). All experiments were conducted in accordance to the guidelines from the Canadian Council on Animal Care in Science, approved by Queen’s University, and were in accordance with the National Institutes of Health guidelines for animal research with the approval of the Institutional Animal Care and Use Committee at the University of North Carolina at Chapel Hill.

### Stereotaxic injections

Adult mice (>8 weeks) were deeply anesthetized with 5% isoflurane (vol/vol) in oxygen, placed into a stereotactic frame (Kopf Instruments, Tujunga, CA, USA) and maintained at 1.5–2.5% isoflurane during surgery. A hole was drilled in the skull using CeA coordinates ML: ±2,95, AP: –1.15, DV: –4.75 from Bregma. Microinjections of 300 nL of virus (AAV5-EF1a-DIO-ChR2-mCherry or AAV5-EF1a-DIO-ChR2-eYFP) were made bilaterally using a 1□μl Neuros Hamilton syringe (Hamilton, Reno, NV, USA) at a rate of 100□nl□/minute. After infusion, the needle was left in place for at least an additional 5□minutes to allow complete diffusion of the virus before being slowly withdrawn. After surgery, all mice were returned to group housing, and recovered for at least 6 weeks prior to the start of experiments.

### Slices preparation and electrophysiology

Mice were anesthetized with isoflurane (5% at 5 L/minute) and their brain removed into ice-cold artificial cerebral spinal fluid (aCSF) containing (in mM): 126 NaCl, 2.5 KCl, 1.2 MgCl2, 6 CaCl2, 1.2 NaH2PO4, 25 NaHCO3, and 12.5 D-glucose equilibrated with 95% O2/5% CO2. Brains were cut in 2°C aCSF into coronal slices (300 μm) with a vibrating blade microtome (VT-1000; Leica Canada, Concord, ON, Canada). The ovBNST slice of interest for this study corresponded to –0.26mm from Bregma. Slices were incubated at 34°C for 60 minutes and transferred to a chamber perfused (2–3 mls/minute) with aCSF at 34°C. Remaining slices were kept in aCSF at room temperature until further use. Whole-cell voltage-clamp recordings were made using glass microelectrodes (3–5 MΩ) filled with (in mM): 70 K+-gluconate, 80 KCl, 1 EGTA, 5 HEPES, 2 MgATP, 0.3 GTP, and 1 P-creatine. Electrical stimuli (10–100 μA, 0.1 ms duration) or optical stimuli (490 nm LED intensity 2–100%, 0.1 ms duration) were applied at 0.1 Hz. Inhibitory post-synaptic currents (IPSCs) were evoked by local fiber stimulation with tungsten bipolar electrodes or by a 490 nm LED via the objective while neurons were voltage-clamped at −70 mV. GABA A-IPSCs were pharmacologically isolated with 6,7-dinitroquinoxaline-2,3-dione (DNQX, 50 µM). To induce local endogenous neuropeptide release, post-synaptic neurons were repetitively depolarized in voltage clamp from –70 to 0 mV (100 ms) at a frequency of 2 Hz for 5 minutes (Normandeau *et al*, 2018). We quantified peak amplitude and defined 3 possible outcomes to postsynaptic depolarization or drug bath application: 1-long-term potentiation (LTP^GABA^; >20% deviation from baseline after 20 minutes), 2-long-term depression (LTD^GABA^; <20% deviation from baseline after 20 minutes) or 3-no change (NC, within 20% deviation from baseline after 20 minutes). Recordings were made using a Multiclamp 700B amplifier and a Digidata 1440A (Molecular Devices LLC, San Jose, CA, USA). Data were acquired and analyzed with Axograph X running on Apple computers and Clampfit on Windows computers.

### Drugs

Stock solutions of SR142948 (10 mM), Norbinaltorphimine (Nor-BNI, 100 mM), NT (1mM) and JMV431 (1mM) were prepared in distilled water. Stock solutions of DNQX (100 mM), and NTRC844 (1mM) were prepared in DMSO (100%). All drugs were further dissolved in the physiological solutions at the desired concentrations (DNQX 50μM, SR142948 10 μM, Nor-BNI, 100 nM, JMV431 100nM, NTRC844 100nM, SR48692 1μM, NT 1μM) and the final DMSO concentration never exceeded 0.1%.

### Fluorescence *In Situ* Hybridization (FISH)

Immediately after removal, brains were placed on a square of aluminum foil on dry ice to freeze for 5 minutes before wrapping to prevent tissue damage. Brains were then placed in a −80□°C freezer for no more than 1 week before slicing. In all, 12- μm slices containing the CeA and ovBNST were obtained on a Leica CM3050S cryostat (Leica Biosystems, Wetzlar, Germany) and placed directly on coverslips. FISH was performed using the Affymetrix ViewRNA 2-Plex Tissue Assay Kit with custom probes for *Nts, Ntsr1*, and *Ntsr2* designed by Affymetrix (Santa Clara, CA, USA). FISH was also done using the Advanced Cell Diagnostics (ACD) HybEZ(TM) II Hybridization System with custom probes for *Nts* and *PDyn* designed by ACD (Newark, CA, USA). Slides were coverslipped with SouthernBiotech DAPI Fluoromount-G. (Birmingham, AL, USA). z-Stack (3 × 5 tiled; 8 optical sections comprising 10.57□μm in total) were obtained on a Zeiss 800 confocal microscope. All images were preprocessed with stitching and maximum intensity projection. Quantification of probe colocalization was hand counted using the cell counter plugin in FIJI (ImageJ, NIH, Bethesda, MD, USA). For all studies, cells were classified into three groups: probe 1+, probe 2+ or probe 1+ and 2+. Only cells positive for a probe were considered.

### Statistical analyses

Changes in IPSCs peak amplitude were measured from baseline and are shown as percentages as follows: (Peakamplitude_post_−Peakamplitude_baseline_/Peakamplitude_baseline_)*100. Data are reported as means ± SEM and each data point represents the average of values in 1-minute bins (6 evoked IPSCs) across recorded neurons. ANOVAs were used to compare multiple means and the appropriate post-hoc statistical tests for multiple comparisons conducted when ANOVAs deemed significance. All statistics from the within-subjects ANOVAs were reported using the Greenhouse-Guesser correction for violations in sphericity, and degrees of freedom for Greenhouse-Giesser values were rounded up to the nearest whole number. A Bonferroni correction was used for multiple comparisons. Fisher’s exact tests analyzed contingency tables of the neuronal response distribution. All statistical analyses were done with SPSS Statistics Version 23 (SAS Institute) or Prism 6 (Graph Pad).

## Results

Multiple neuropeptides are usually co-represented in individual neurons in the brain, suggesting concerted roles in coordinating proper neural circuit function. However, whether and how multiple neuropeptidergic systems interact neurophysiologically to regulate neural circuits is largely unknown since neuropeptides are often studied in isolation. Here, we triggered release of endogenous neuropeptides in brain slices to investigate how neuropeptides can regulate CeA to ovBNST inhibitory circuit in male mice.

### Bi-directional modulation of electrically-evoked GABA-IPSCs by NT and Dyn in the ovBNST of male mice

Electrical stimulation of local fibers evoked GABA_A_-IPSCs (eIPSCs) in ovBNST neurons and peak amplitudes were measured. Upon stable baseline of eIPSCs (minimum 5 minutes), post-synaptic ovBNST neurons were activated for 5 minutes by repetitive depolarization (0mV, 100msec, 2Hz), which resulted in long-term potentiation (LTP^GABA^) of eIPSCs in 90% of recorded neurons (time x group, F_3,8_=9.5, p=0.002; Figure 1B). The addition of non-selective NTRs antagonist (SR-142948, 10μM) diminished LTP^GABA^ cell response indicating that NT acts as an important modulator of inhibitory transmission in mice (Fisher’s exact test (aCSF vs. SR); p=0.04; Figure 1C, E). Blocking NT receptors also uncovered LTD^GABA^ that was ablated by the co-application of the selective KOR antagonist nor-NBI (100nM) with SR (time x group, F_1,9_=31.6, p=0.0003; Figure 1D, E). As such, locally released NT and Dyn inversely regulate inhibitory transmission in the ovBNST.

**Figure 1:**
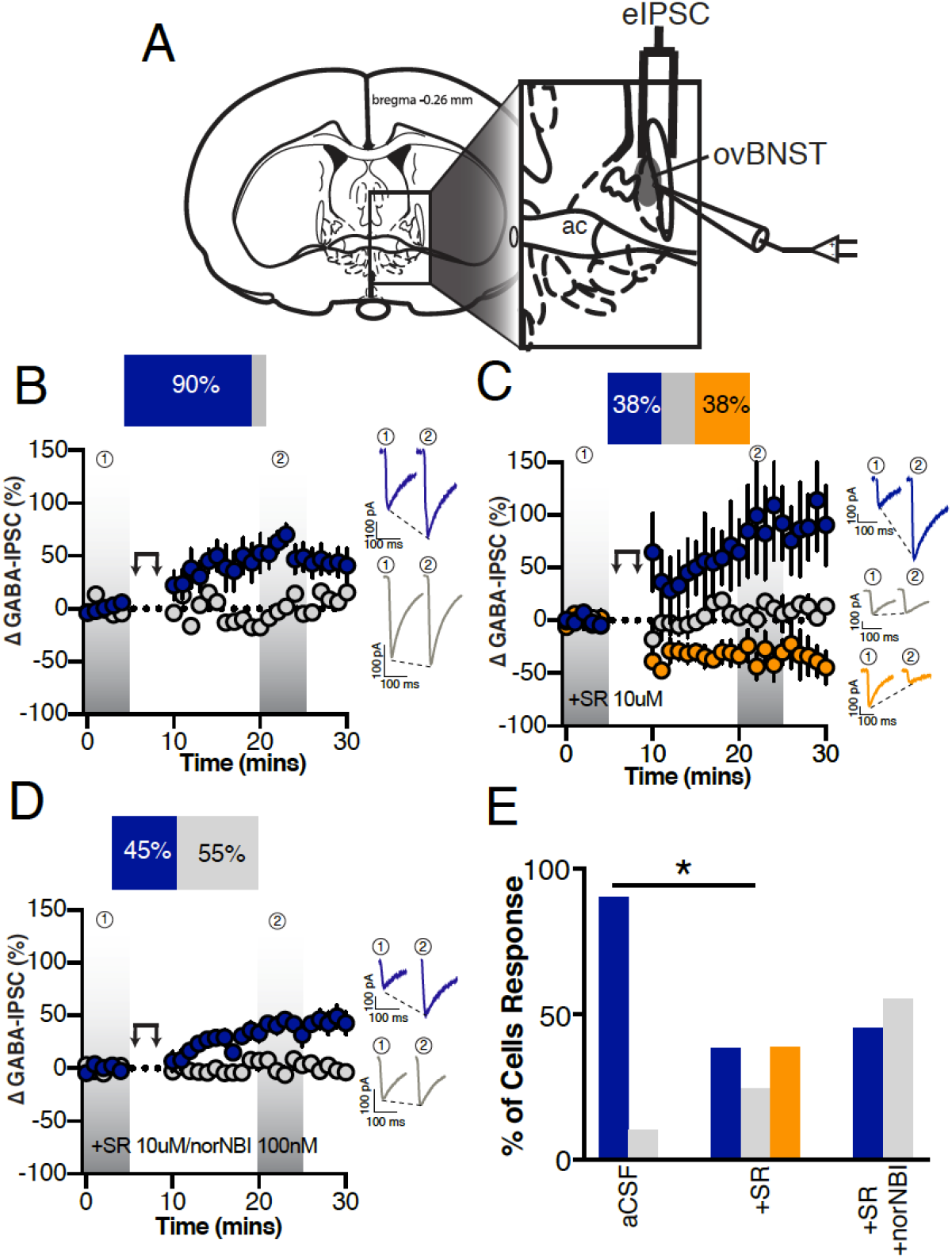
Endogenous neuropeptides modulation of electrically-evoked ovBNST GABA_A_ IPSCs. A, Schematic illustrating stimulating and recording electrodes placement in mice brain slices containing the ovBNST. Recordings were restricted to the displayed shaded oval area. B-D, Effects of postsynaptic depolarization (double arrow symbol) on binned (1 minute, 6 events) electrically-evoked GABA_A_-IPSCs in (B) aCSF (n=10 cells/6 mice), (C) the presence of the non-selective NTR antagonist SR142948 (10μM, n=8 cells/5 mice) or (D) SR142948 + the KOR antagonist norNBI (100nM, n=11 cells/4 mice). Insets in B-D are representative electrically-evoked GABA_A_-IPSCs before and after postsynaptic depolarization (double arrows). E, Histogram summarizing the proportion of responding neurons to post-synaptic activation across different pharmacological treatments. Blue LTP, grey no change and orange LTD. Asterisks, p<0.05.

### Locally released NT and Dyn target CeA-to-ovBNST GABA-IPSCs

Next, using V*gat-Cre* male mice injected with DIO-ChR2-eYFP/mCherry in the CeA, we light-activated (LED 490nm) specific CeA inputs and evoked optical V*gat*^CeA□ovBNST^ GABA-IPSCs (opIPSCs) in ovBNST neurons. Post-synaptic activation of ovBNST neurons resulted in 54% LTP^GABA^ of opIPSCs and 8% LTD^GABA^ (time x group, F_5,20_=4.1, p=0.01; Figure 2C). Application of the non-selective NTRs antagonist robustly reduced LTP^GABA^(8%) and revealed further LTD^GABA^ (42%) (Fisher’s exact test (aCSF vs. SR), p=0.04; Figure 2D, F). Combined application of NTRs and KOR antagonist completely eliminated LTP^GABA^ and LTD^GABA^ cell responses (Fisher’s exact test (SR vs. SR/norNBI), p=0.04; Figure 2E, F). Therefore, NT and Dyn are dominant modulators of CeA inputs onto ovBNST in mice.

**Figure 2:**
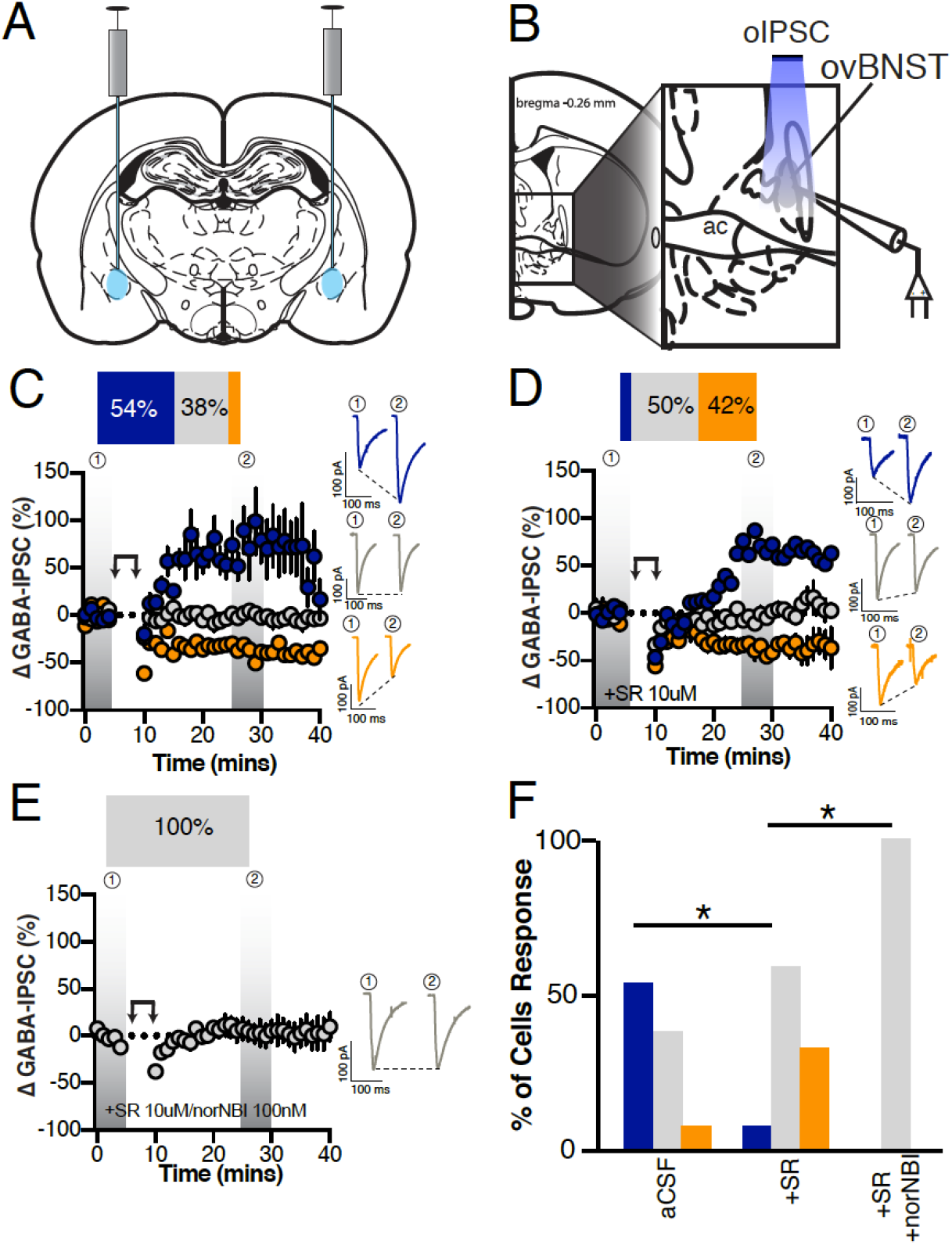
Effect of postsynaptic depolarization on optically-evoked V*gat*^CeA→ovBNST^ GABA_A_ IPSCs. A, Illustration demonstrating ChR2 bi-lateral injections into the CeA of *Vgat-Cre* mice. B, Illustration demonstrating optically stimulated recordings of ovBNST neurons. C-E, Post-synaptic depolarization in male V*gat-Cre* mice injected with ChR2 in the CeA and recorded from ovBNST brain slices in (C) aCSF (n=13 cells/6 mice), (D) in the presence of the non-selective NTR antagonist SR142948 (10μM, n=12 cells/5 mice), (E) SR142948 + the KOR antagonist norNBI (100nM, n=8 cells/2 mice). Insets in C-E are representative optically-evoked GABA_A_IPSCs before and after postsynaptic depolarization (double arrows). F, Histogram summarizing the proportion of responding neurons to post-synaptic activation across different pharmacological treatments. Blue LTP, grey no change and orange LTD. Asterisks, p<0.05.

### Expression of *Nts* and *Pdyn* mRNA co-localized in the ovBNST

Because NT- and Dyn-mediated modulation of inhibitory transmission was detectable in a vast majority of ovBNST neurons, we hypothesized that both neuropeptides might be highly co-localized. Dual fluorescent *in situ* hybridization (FISH) revealed co-localization of preprodynorphin (*Pdyn*) and NT (*Nts*) mRNA in ovBNST neurons (66% of neurons, Figure 3).

**Figure 3:**
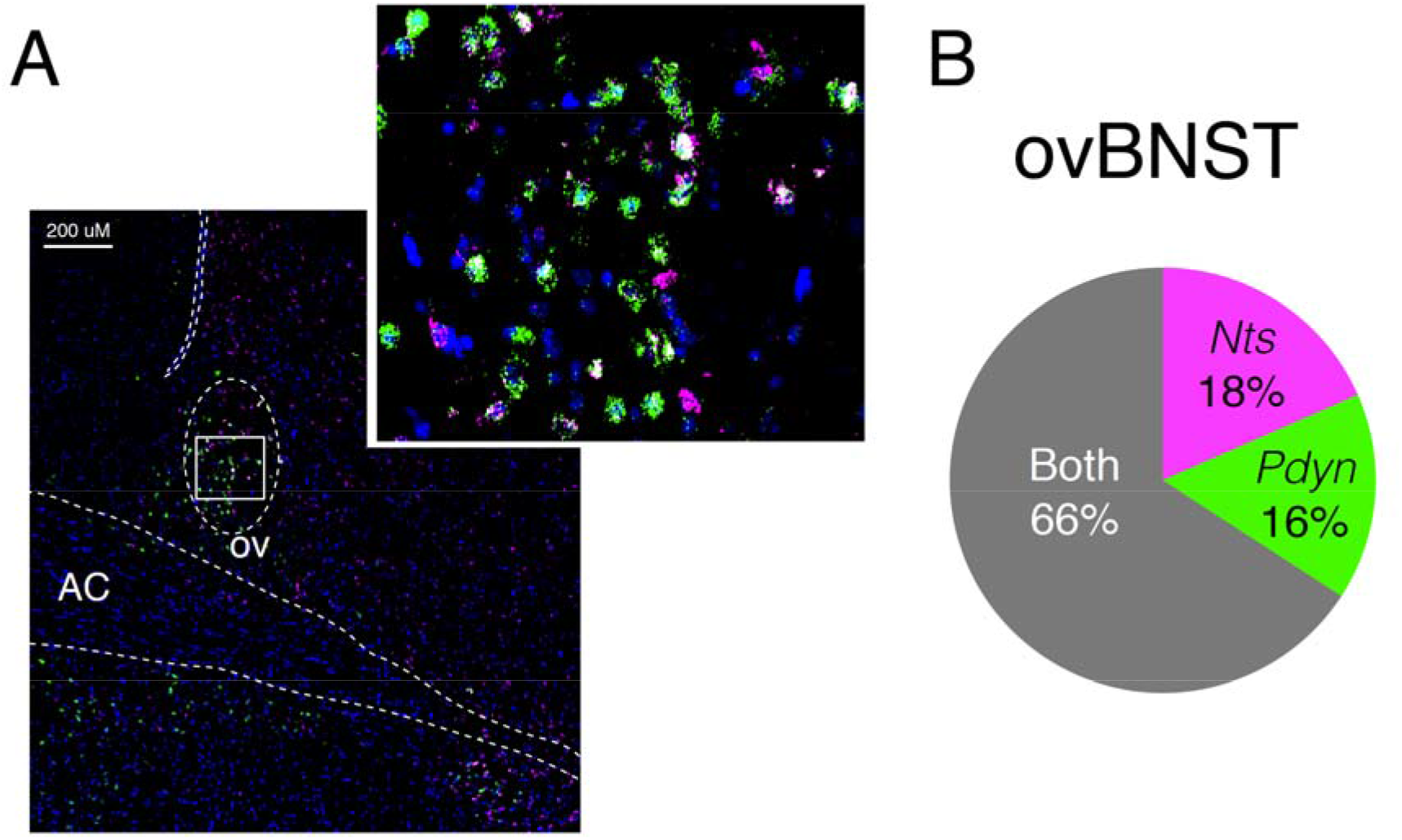
Expression and co-localization of *Pdyn* and *Nts* mRNA in the ovBNST. A, Representative image of dual fluorescent *in situ* hybridization for *Nts/Pdyn* (*Nts* (green), *Pdyn* (purple), and DAPI (blue)). B, Distribution of *Nts, Pdyn* and co-co-localizing (both) mRNA expressing cells (n=4 mice, 4 slices/mouse).

### NTR1 and NTR2 bi-directionally modulated electrically evoked GABA-IPSCs in the ovBNST

Lastly, NT binds to two different G-protein receptors: NTR1 and NTR2 (Vincent *et al*, 1999). Both *Ntsr1* and *Ntsr2* mRNA were present in the ovBNST and CeA (Figure 4A-H). However, neither *Ntsr1* nor *Ntsr2* co-localized extensively with *Nts* mRNA suggesting that NT acted pre-synaptically or on another population of *Ntsr*-positive neurons in the ovBNST.

**Figure 4:**
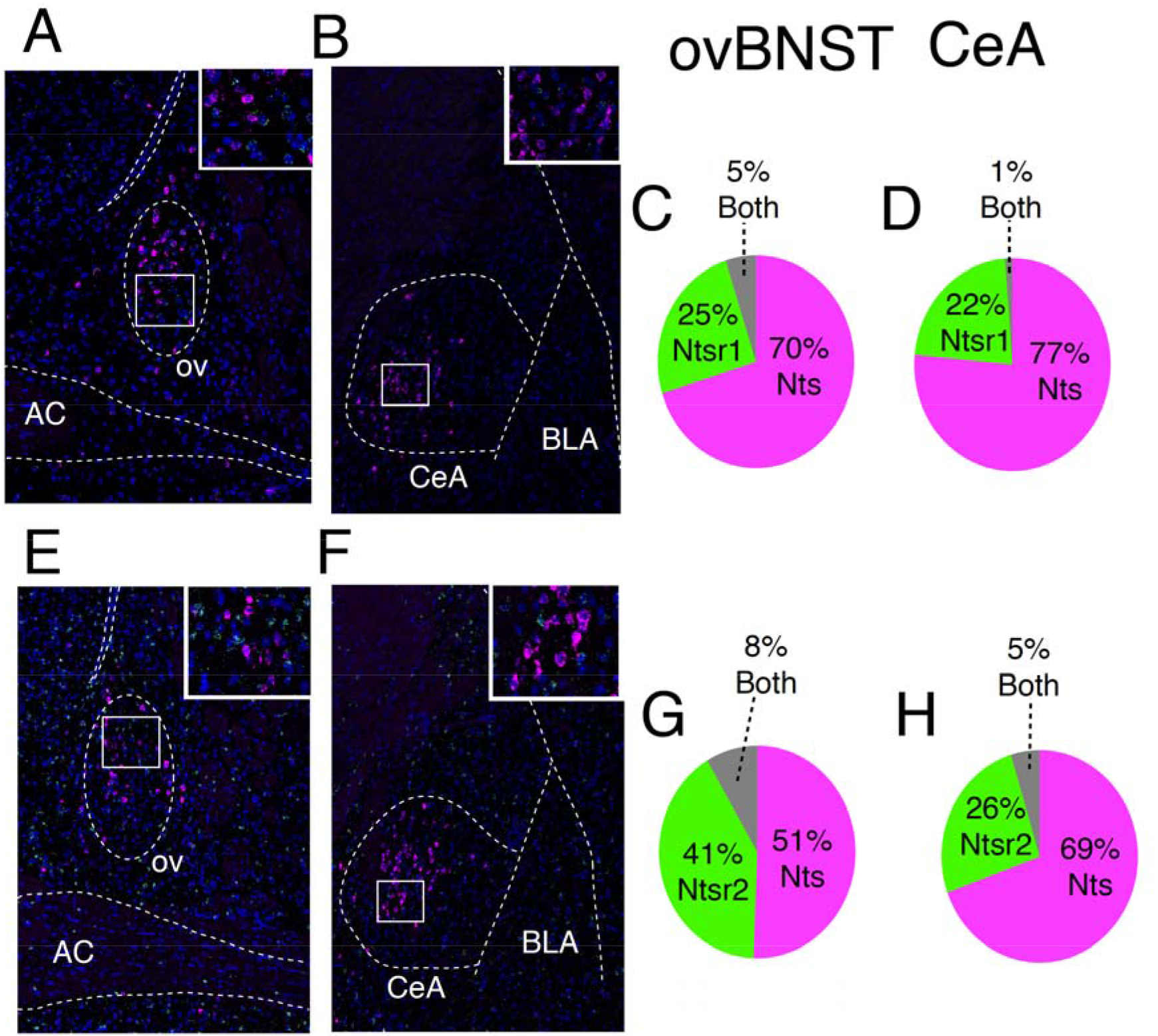
Expression and co-localization of NT and NTRs mRNA in the ovBNST and CeA. Representative image of dual fluorescent *in situ* hybridization (FISH) for *Nts/Ntsr1* (*Nts* (purple), *Ntsr1* (green), DAPI (blue) in the ovBNST (A) and CeA (B). Distribution of *Nts, Ntsr1*and co-localizing mRNA expression cells (n=2 mice, 2 slices/mouse) in the ovBNST (C) and CeA (D). Representative image of dual FISH for *Nts/Ntsr2* (*Nts* (purple), *Ntsr2* (green), DAPI (blue)) in the ovBNST (E) and CeA (F). Distribution of *Nts, Ntsr2*and co-co-localizing (both) mRNA expressing cells (n=2 mice, 2 slices/mouse) in the ovBNST (G) and CeA (H).

Pharmacological application of NT (1 μM) for 5 minutes resulted in LTP^GABA^ of eIPSCs in 80% of recorded neurons (time, F_1,46_=8.3, p=0.002; Figure 5A). Bath application of the NTR2 selective antagonist (NTRC844, 100nM) did not significantly alter cell responses to bath-applied NT with 57% of cells responding with LTP^GABA^ (Fisher’s exact test (NT vs. NT+NTRC844), p=0.6; Figure 5B, E) (Thomas *et al*, 2016). Bath application of the NTR1 selective antagonist (SR48692, 1μM) however, reduced the percentage of neurons responding with LTP^GABA^ (Fisher’s exact test (NT vs. NT +SR48692), p=0.002; Figure 5C, E), suggesting that NTR1 are required to mediate the post-synaptic activation induced by NT. NTR1 blockade also uncovered LTD^GABA^ in 80% of neurons that could be NTR2 mediated. Accordingly, bath application of the NTR2 selective agonist (JMV431, 100nM) for 5 minutes resulted in a LTD^GABA^ in 38% of neurons (time x group, F_6,35_=3.4, p=0.04; Figure 5D) (Thomas *et al*, 2014). Overall, NTRs have opposing modulatory roles on inhibitory transmission in the ovBNST, although the NTR1 component was predominant in response to endogenous NT release.

**Figure 5:**
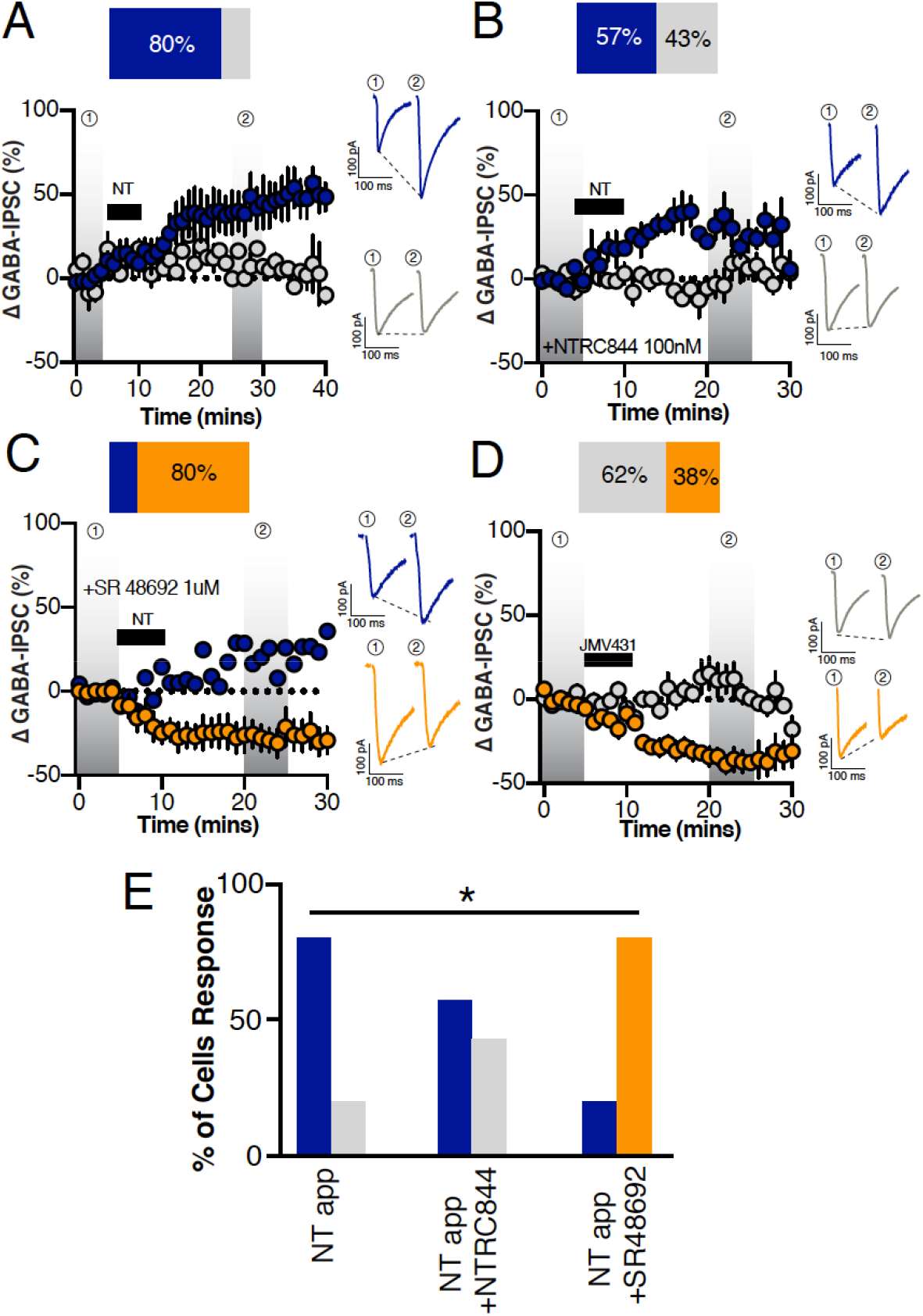
Contribution of neurotensin receptors 1 and 2 on exogenous NT-induced modulation of electrically-evoked ovBNST GABA_A_-IPSCs. A, Effect of a 5-minute bath application of NT (1uM) (black horizontal bar) on the peak amplitude of eIPSCs (n=8 cells/4 mice). Effect of NT (1µM, black horizontal bar) on eIPSCs in the presence of (B) the NTR2-selective antagonist NTRC844 (100 nM, n=7 cells/3 mice) or the (C) NTR1-selective antagonist SR48692 (1 μM, n=5 cells/3 mice) D, Effect of a 5-minute bath application of NTR2 agonist JMV431 (100nM, n=8 cells/4 mice) (black horizontal bar) on the peak amplitude of eIPSCs in the ovBNST. Insets in B-D are representative electrically-evoked GABA_A_-IPSCs before and after bath application of NT or JMV431. Blue LTP, grey no change and orange LTD. Asterisks, p<0.05.

## Discussion

Post-synaptic depolarization of ovBNST neurons in male mice brain slices resulted in the putative release of the neuropeptides NT and Dyn, respectively increasing and decreasing inhibitory inputs originating in the CeA. FISH revealed that Nts and *Pdyn* mRNA significantly overlapped in single ovBNST cells demonstrating the co-localization of NT and Dyn in the ovBNST. Locally released NT and Dyn modulated CeA inhibition of ovBNST neurons through NTR1 and KOR, respectively. Together, our data shed new light unto the coordinated action of co-localized neuropeptides in fine-tuning neural circuit function.

Postsynaptic activation of voltage-clamped ovBNST neurons resulted in a robust and long-lasting modulation of inhibitory postsynaptic GABA_A_ currents in a vast majority (up to 90%) of recorded neurons. We hypothesized that these could be neuropeptides since they are generally released after high or prolonged postsynaptic depolarization, and have long-lasting effects (Karhunen *et al*, 2001; Whim and Lloyd, 1989). Pharmacological blockade of NTR and KOR receptors confirmed that NT and Dyn were fully responsible for the bi-directional regulation of ovBNST inhibitory inputs from the CeA. In male rats, the same post-synaptic activation protocol (2Hz, 5 minutes) also triggered local endogenous neuropeptides release and similar modulation of inhibitory synaptic transmission in the ovBNST, suggesting a conserved mechanism across both species (Normandeau *et al*, 2018). Although our data do not preclude the presence of other neuromodulators (for example endocannabinoids, growth factors, or other neuropeptides), the modulation of inhibitory synaptic transmission was largely eliminated by the co-application of NTR and KOR antagonists. Thus, these observed effects were predominantly NT and Dyn dependent.

Postsynaptic activation resulted in a strikingly homogenous NT-mediated enhancement of inhibitory transmission in the ovBNST. This is consistent with the prevalence of NT-positive neurons in the ovBNST (Allen Mouse Brain Atlas, Ju *et al*, 1989b). This homogenous neurophysiological outcome we measured here in mice and previously in rats, may seem to contrast with the various neurophysiological signatures reported in the rat dorsal BNST neurons (Hammack 2007, Dabrowska, 2013). But yet, a majority of dorsal BNST neurons express the neuropeptide CRF and further co-express NT regardless of their neurophysiological signature, perhaps explaining the homogenous neurophysiological outcome we measured upon postsynaptic depolarization (Dabrowska *et al*, 2013; Hammack *et al*, 2007; Ju and Han, 1989a). Importantly, we purposefully restricted our recordings to the oval subregion of the dorsolateral BNST (see A and Figure 2B) and that may have enabled us to target a population of neurons. Nonetheless, the majority of neurons in this sub-nuclei are similar morphologically and we suspect the ovBNST may function in a coordinated network fashion (Larriva-Sahd, 2006).

Narrowing our investigation to CeA-mediated optically-driven (op)IPSCs revealed that post-synaptic depolarization correspondingly resulted in robust modulation of inhibitory inputs onto ovBNST neurons. Although at a lower proportion than non-specific (eIPSC) input stimulation, a majority of neurons (54%) displayed LTP^GABA^ of similar magnitude and duration. In contrast, LTD^GABA^ was more readily observable with opIPSCs (8% of neurons) without NTRs blockade. Similar to eIPSC, NTR blockade fully revealed KOR-mediated LTD^GABA^. Co-application of NTR and KOR antagonists completely abolished the modulation of opIPSCs whereas a residual LTP^GABA^ remained with electrical stimulation (45%). In male rats, a similar residual eIPSC LTP^GABA^ was mediated by CRF, and we suspect this neuropeptide may also be responsible for the residual LTP^GABA^ observed in mice (Normandeau *et al*, 2018). Nonetheless, our findings strongly suggest that NT and Dyn are fully responsible for bi-directional modulation of CeA inhibitory inputs to ovBNST neurons in male mice (Li *et al*, 2012). Both *Pdyn* and *Nts* mRNA were highly co-localized in the ovBNST suggesting that they could be released from the same cells to have their modulatory effects. In contrast, *Kor* mRNA is only expressed in the CeA while *Ntsr1* and *Ntsr2* mRNA were expressed in both ovBNST and CeA neurons (Poulin *et al*, 2009; Torruella-Suarez, 2018). Neither neuropeptides mRNA (NT or Dyn) co-localized significantly with their respective receptors mRNA (Torruella-Suarez, 2018). Therefore, we suggest that NT and Dyn found in ovBNST neurons act directly onto pre-synaptic axon terminals originating from the CeA (illustrated in Figure 6). Though, we cannot exclude the possibility that NT may be working through interneurons since we identified *Ntsr* mRNA in the ovBNST.

**Figure 6:**
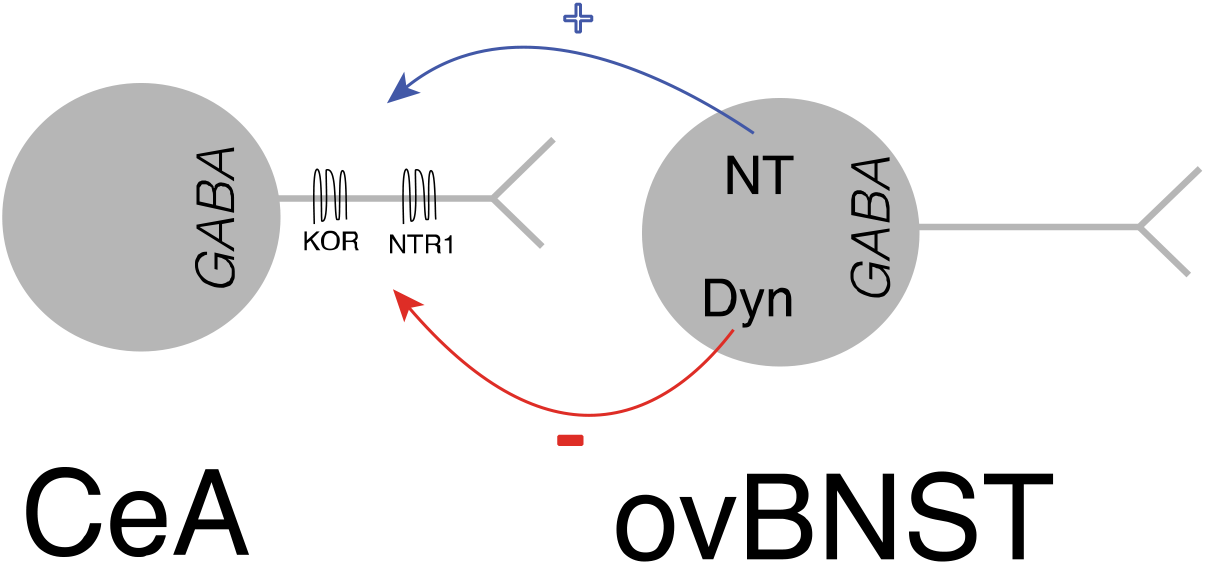
Illustration of NT and Dyn bi-directional modulation of CeA inputs onto ovBNST neurons.

Bath-application of exogenous NT resulted in LTP^GABA^ in 80% of recorded ovBNST neurons, similar to our previous observation in rats (Krawczyk *et al*, 2013). NTR1 (SR48692), but not NTR2 (NTRC844), blockade, significantly interfered with NT-induced LTP^GABA^. Interestingly, the NTR2 specific agonist JMV431 resulted in LTD^GABA^ in 38% of recorded neurons, indicating that both NT receptors may have opposing modulatory roles on inhibitory transmission in the ovBNST. NT has a higher affinity to NTR1 than NTR2, suggesting that post-synaptic activation may release low to moderate amounts of endogenous NT that may only activate NTR1s (Tschumi and Beckstead, 2018). Additionally, NTR2 are preferentially expressed intra-cellularly whereas NTR1s are expressed at the cell membrane, potentially increasing NT’s ability to bind and activate NTR1s (Perron *et al*, 2007).

NT and Dyn may independently modulate inhibition in the ovBNST by acting at their respective receptors. However, they may also interact directly since KOR and NTR1 can form heterodimers that alters KOR-mediated signaling (Liu *et al*, 2016). In fact, KOR and NTR1 are both expressed in the CeA and could co-localize pre-synaptically in ovBNST inputs (Boudin *et al*, 1996; Mansour *et al*, 1987). While the exact functional link between NT and Dyn remains elusive, it is clear that the two neuropeptides have strong cooperative, although antagonistic, potential to regulate inhibition in the ovBNST.

With either stimulation of local fibers or CeA inputs, the blockade of NTR1 uncovered Dyn-induced LTD^GABA^. Consequently, NT seems to be the primary modulator of CeA inhibitory inputs onto ovBNST neurons. Our stimulation paradigm (2Hz, 5minutes) could particularly favor NT over Dyn release, as previous research suggests that different firing patterns can release different neuropeptides (Poulain *et al*, 1977). Alternatively, the relative availability of neuropeptides and receptor expression or function may be state-dependent. Here, the mice were sated, minimally stressed, and brain slices prepared approximately 2 hours within the active (dark) phase of the mice’s circadian cycle. Because the ovBNST seems important for energy homeostasis and is a one of the brain’s circadian clock, it will be critical to determine whether Dyn-mediated LTD^GABA^ may not become the primary response in different conditions (Amir *et al*, 2004; Dong *et al*, 2001).

Anatomical studies suggest that the CeA and ovBNST robustly inhibit one another such that their physiological roles are possibly antagonistic (Dong *et al*, 2001; Petrovich *et al*, 1997). The CeA is instrumental in promoting aversive learning and fear response whereas the dorsolateral BNST might be more related to appetitive behaviours (Davis *et al*, 2010; Jennings *et al*, 2013). Energy homeostasis may be an important factor in the neuromodulatory effects of NT vs. Dyn on the CeA-to-ovBNST circuit. For instance, they have opposing effects on foraging behaviours: NT and Dyn respectively decrease and increase food intake in both rats and mice (Cooke *et al*, 2009; Lambert *et al*, 1993; Levine *et al*, 1983; Luttinger *et al*, 1982; Sainsbury *et al*, 2007). Intriguingly, NT reduces Dyn-induced feeding in rats (Levine *et al*, 1983). Furthermore, NT and Dyn expression and release are metabolic state-dependent. In rats, NT release into the hepatic-portal circulation occurs immediately after cessation of eating (George *et al*, 1987). In contrast, Dyn expression increases significantly in the rat hypothalamus after a 72-hour food deprivation (Przewlocki *et al*, 1983). Therefore, NT may promote CeA inhibition of the ovBNST to decrease foraging behaviour.

In sum, our data strongly suggest that NT and Dyn cooperate to fine-tune CeA inhibitory inputs onto ovBNST neurons. We have described one circuit, as well as one pair of behaviorally-relevant neuropeptides that could serve as a source of bi-directional modulation *in vivo*: the physiological and/or behavioural relevance of this mechanism remains to be elucidated. Our study paves the way to investigate whether this phenomenon occurs in other neuropeptidergic systems, or can be generalized outside of the extended amygdala.

